# The endoplasmic reticulum-plasma membrane tethering protein TMEM24 is a regulator of cellular Ca^2+^ homeostasis

**DOI:** 10.1101/2021.06.23.449694

**Authors:** Beichen Xie, Styliani Panagiotou, Jing Cen, Patrick Gilon, Peter Bergsten, Olof Idevall-Hagren

## Abstract

Endoplasmic reticulum (ER) - plasma membrane (PM) contacts are sites of lipid exchange and Ca^2+^ transport, and both lipid transport proteins and Ca^2+^ channels specifically accumulate at these locations. In pancreatic β-cells, both lipid- and Ca^2+^ signaling are essential for insulin secretion. The recently characterized lipid transfer protein TMEM24 dynamically localize to ER-PM contact sites and provide phosphatidylinositol, a precursor of PI(4)P and PI(4,5)P_2_, to the plasma membrane. β-cells lacking TMEM24 exhibit markedly suppressed glucose-induced Ca^2+^ oscillations and insulin secretion but the underlying mechanism is not known. We now show that TMEM24 only weakly interact with the PM, and dissociates in response to both diacylglycerol and nanomolar elevations of cytosolic Ca^2+^. Release of TMEM24 into the bulk ER membrane also enables direct interactions with mitochondria, and we report that loss of TMEM24 results in excessive accumulation of Ca^2+^ in both the ER and mitochondria and in impaired mitochondria function.

## INTRODUCTION

Lipid exchange between the endoplasmic reticulum (ER) and the plasma membrane (PM) is facilitated by lipid-transport proteins that are concentrated to, and participate in the formation of, ER-PM contact sites. These junctions are also important sites for cellular Ca^2+^ homeostasis, and the lipid transport is often coupled to changes in the cytosolic Ca^2+^ concentration (Balla, 2018; Chung et al., 2017; Saheki & De Camilli, 2017). The changes in membrane lipid concentrations occurring as a consequence of lipid transport may also influence Ca^2+^ influx or extrusion through modulation of Ca^2+^ channel activity or clustering (Johnson et al., 2018; Suh et al., 2010; Sun et al., 2019; Xie et al., 2016). TMEM24 is a recently characterized lipid transport protein that localize to ER-PM contacts where it, through an N-terminal synaptotagmin-like mitochondrial-lipid-binding (SMP) domain, provides the plasma membrane with phosphatidylinositol, the precursor of the signaling lipids PI(4)P and PI(4,5)P_2_ (Lees et al., 2017; Sun et al., 2019). It binds negatively charged lipids in the plasma membrane via a C-terminal polybasic region, and neutralization of positively charged amino acids in this region by PKC-dependent phosphorylation results in TMEM24 dissociation from the plasma membrane. TMEM24 is also equipped with a C2-domain, which, however, seems dispensable for both plasma membrane binding and lipid transport. The spatial regulation of TMEM24 activity differs from other SMP- and C2-domain proteins, such as the Extended synaptotagmins, whose interaction with the plasma membrane and lipid transport are instead triggered by Ca^2+^ elevation (Bian et al., 2018; Giordano et al., 2013; Idevall-Hagren et al., 2015).

Pancreatic beta-cells are secretory cells that produce insulin and release this hormone to the circulation in response to elevated blood glucose levels. The mechanism controlling insulin secretion is well-characterized and involves glucose uptake and metabolism, resulting in an elevated ATP/ADP ratio which in turn closes ATP-sensitive K^+^ (K_ATP_)-channels, causing membrane depolarization, opening of voltage-dependent Ca^2+^ channels, Ca^2+^ influx and the fusion of insulin-containing granules with the plasma membrane (Rorsman & Ashcroft, 2018). Ca^2+^ is the most important trigger of insulin secretion, but the response can also be modulated by many factors, including lipids and lipid-derived signaling molecules. Phosphoinositides in particular has been shown to regulate insulin secretion at several stages, including membrane depolarization, Ca^2+^ influx and granule docking and release (Wuttke, 2015). The phosphoinositide PI(4,5)P_2_ also serves as a precursor of inositol 1,4,5-trisphosphate (IP3), which triggers Ca^2+^ release from the ER and may contribute to insulin granule exocytosis, and diacylglycerol (DAG), which together with Ca^2+^ amplifies secretion by stimulating the protein kinase C activity (Wuttke et al., 2013, 2016). Lipid transport at ER-PM contact sites are important for the normal function of insulin-secreting β-cells. DAG transport by E-Syt1 was recently found to provide negative feedback on insulin secretion after being recruited to sites of Ca^2+^ influx where it locally clears the plasma membrane of the pro-secretory lipid (Xie et al., 2019). β-cells with reduced E-Syt1 expression thus exhibits increased accumulation of plasma membrane DAG and excess insulin secretion in response to glucose. E-Syt1 occupies the same contact sites as TMEM24. However, their presence at these contacts does not overlap in time due to their inverse dependence of plasma membrane binding on the prevailing Ca^2+^ concentration (Xie et al., 2019). Interestingly, TMEM24 was recently found to be indispensable for glucose-stimulated insulin secretion (Lees et al., 2017; Pottekat et al., 2013). The complete loss of insulin secretion in clonal TMEM24 knockout β-cells is likely caused by an effect on β-cell Ca^2+^ homeostasis. β-cells lacking TMEM24 thus exhibit markedly suppressed glucose-induced Ca^2+^ oscillations, but the mechanism behind this is not clear (Lees et al., 2017). One possibility is that TMEM24 is necessary for maintaining plasma membrane PI(4,5)P_2_ required for ion channel gating during stimulation, although the global PI(4,5)P_2_ levels are unaltered in TMEM24 KO cells (Lees et al., 2017). The dynamics of TMEM24 in glucose-stimulated β-cells is also difficult to reconcile with a role as a positive regulator of insulin secretion, since it is spatially separated from this process during glucose stimulation. Clarification of the role of TMEM24 in the regulation of insulin secretion is therefore required.

We now show that the subcellular distribution of TMEM24 is highly dynamic and influenced by both Ca^2+^ and plasma membrane DAG. In contrast to previous studies, we do not find a requirement of TMEM24 for normal Ca^2+^-triggered insulin secretion. Instead, we identified TMEM24 as an important regulator of intracellular Ca^2+^ stores, and show that β-cells lacking TMEM24 exhibit both ER and mitochondrial Ca^2+^ overloading, resulting in dissipation of the mitochondrial membrane potential and in impaired oxidative phosphorylation.

## MATERIALS AND METHODS

### Plasmids and Reagents

TMEM24-EGFP (Lees et al., 2017), E-Syt1-GFP (Giordano et al., 2013) and mRFP-PH-PLCδ1 were gifts from Pietro De Camilli (Yale University). VAMP2-pHluorin and NPY-mCherry were gifts from Sebastian Barg (Uppsala University). GFP-P4M-SidM was a gift from Gerald Hammond (University of Pittsburgh, Addgene plasmid 51472)(Hammond et al., 2014). GFP-C1aC1b_PKC_ was a gift from Anders Tengholm (Uppsala University)(Wuttke et al., 2013). R-GECO and mito-LAR-GECO were gifts from Robert Campbell (University of Alberta, Addgene plasmid 32444 and 61245) (Wu et al., 2014; Zhao et al., 2011). mApple-Tomm20 was a gift from Michel Davidson (Addgene plamid 54955). TMEM24-mCherry was generated by PCR amplification of human TMEM24 with flanking Nhe1 and EcoR1 followed by ligation into the mCherry-N1 vector using the following primers: TMEM24-Nhe1-fwd, CTAGCTAGCATGGATCCGGGCTGGGGGCA; TMEM24-EcoR1-rev, CCGGAATTCTGAGCTGGGGGCTGGGGTT. All salts, HEPES, poly-_L_-lysin, EGTA, nitrilotriacetic acid (NTA) and diazoxide were from Sigma-Aldrich. DMEM, Penicillin, streptomycin, glutamine and FBS were from Life technologies. Carbachol, Phorbol 12-Myristate 13-Acetat (PMA), cyanide-p-trifluoromethoxyphenylhydrazone (FCCP), rotenone, antimycin A and cyclopiazonic acid (CPA) were from Sigma-Aldrich. Bafilomycin was from TOCRIS Bioscience. Fluo-4-AM, Fura-2-AM and TMRM were from Life Technologies and Cal-520-AM was from AAT Bioquest.

### Cell culture and transfection for imaging

The mouse β-cell line MIN6 (passages 18-30) (Miyazaki et al., 1990) was cultured in DMEM (Life Technologies) supplemented with 25 mmol/l glucose, 15% FBS, 2 mmol/l L-glutamine, 50 μmol/l 2-mercaptoethanol, 100 U/ml penicillin and 100 μg/ml streptomycin. The cells were kept at 37°C and 5% CO_2_ in a humidified incubator. Prior to imaging, 0.2 million cells were resuspended in 100 μl Opti-MEM-I medium (Life technologies) with 0.2 μg plasmid (total) and 0.5 μl lipofectamine 2000 (Life technologies) and seeded in the centre of a 25-mm poly-L-lysine-coated coverslip. The transfection reaction was terminated after 4-6h by the addition of 2 mL complete culture medium and cells were imaged 18-24 h later.

### Alpha-toxin permeabilization

Transfected MIN6 cells, grown on 25-mm poly-L-lysine-coated glass coverslips, were incubated with 0.5 μM of the AM-ester form of Cal520 for 30 min at 37°C. The coverslips were subsequently used as exchangeable bottoms in a modified Sykes-Moore open superfusion chamber that was mounted on the stage of a TIRF microscope (described below) and connected to a peristaltic pump that allowed rapid change of medium. Following change from normal, extracellular-like, medium (125 mM NaCl, 4.9 mM KCl, 1.3 mM MgCl_2_, 1.2 mM CaCl_2_, 25 mM HEPES, 1 mg/ml BSA with pH set to 7.4) to an intracellular-like medium (see below), the superfusion was interrupted and alpha-toxin was added directly to the chamber (final concentration ≈50 μg/mL). Permeabilization was considered complete when the Cal520 fluorescence had decreased by >90%, which typically took 2-5 min. Superfusion was then started again and the cells were exposed to intracellular-like buffers containing calibrated Ca^2+^ concentrations while fluorescence from both remaining Cal520 and mCherry-tagged fusion proteins was recorded. These experiments were performed at ambient temperature (21-23 °C).

### Intracellular-like media

Intracellular-like media with buffered pH, [Ca^2+^] and [Mg^2+^] used in alpha-toxin permeabilization experiments contained: 6 mM Na^+^, 140 mM K^+^, 1 mM (free) Mg^2+^, 0-100 μM (free) Ca^2+^, 1 mM Mg-ATP, 10 mM HEPES, 2 mM (total) EGTA and 2 mM (total) Nitrilotriacetic acid (NTA) with pH adjusted to 7.00 at 22°C with 2M KOH. The total concentration of Ca^2+^ and Mg^2+^ was calculated using the online version of MaxChelator (http://www.stanford.edu/~cpatton/webmaxcS.htm). Media were made fresh on the day of experiment and kept on ice. To validate the media composition, cells loaded with the Ca^2+^ indicator Cal520 were mounted on a TIRF microscope, permeabilized and exposed to media with increasing Ca^2+^ concentrations. From these data a dose-response curve was generated and the EC_50_ for Ca^2+^-binding to Cal520 was estimated to be 980 nM, an *in situ* estimation 3-fold higher than the reported in vitro K_D_ of 320 nM.

### TMEM24 knockdown by siRNA

MIN6 cells were resuspended with 25 nM antiTMEM24 siRNA smartpool (Dharmacon, siGENOME) pre-mixed with 1.6 μl/ml Lipofectamine RNAiMAX (Life technologies) in Opti-MEM I medium. After 3 hours, the Opti-MEM I medium was replaced by DMEM growth medium containing 25 nM siRNA and 1.6 μl/ml Lipofectamine 3000 (Life technologies) and incubated overnight. Cells transfected with a scrambled sequence siRNA (Dharmacon, siGENOME) following the same procedure were used as controls. After 48 hours, cells were transfected for imaging as described above. The knockdown efficiency was evaluated by western blotting and real-time PCR (see below).

### Generation of TMEM24 knockout MIN6 cells by CRISPR/Cas9

TMEM24 KO MIN6 were generated using a mix of three TMEM24 CRISPR/Cas9 plasmids (Santa Cruz Biotechnology, sc-428047), containing GFP and the following gRNAs: TTACCATGGTGCGCTCTGAT, CAGCACTCAGCCCGCCATGA and GTCCCCCGCTGCCGTCTCCA. TMEM24 CRISPR/Cas9 plasmids were transfected into MIN6 by Lipofectamine 3000 (ThermoFisher, L3000015). After 24-hours, single cells positive (knockout) and negative (control) for GFP were sorted using a BD FACSariaIII CellSorter and subsequently plated into 96-well plates. Cells were cultured in standard MIN6 growth medium. The clones were tested for the presence of TMEM24 by western blot utilizing anti-TMEM24 antibody (ThermoFisher, A304-764A).

### TIRF and confocal microscopy

A medium containing 125 mM NaCl, 4.9 mM KCl, 1.3 mM MgCl_2_, 1.2 mM CaCl_2_, 25 mM HEPES, 1 mg/ml BSA with pH set to 7.4 at 37 °C was used in all microscopy experiments. Cells were pre-incubated in this medium supplemented with 3 mM glucose for 30 min followed by perifusion with the same medium during recordings. A previously described custom-build prism-type TIRF microscope setup equipped with a 16X/0.8-NA water immersion objective (Nikon) was used to observe large population of cells (Idevall-Hagren et al., 2010). It was built around an E600FN upright microscope (Nikon) enclosed in a Perspex box thermostated at 37 °C. 491-nm and 561-nm DPSS lasers (Cobolt, Sweden) were used to excite GFP and mCherry, respectively. The laser beams were merged by dichroic mirrors (Chroma Thechnology) and homogenized by a rotating light shaping diffuser (Physic Optics Corps) before being refocused through a dove prism (Axicon) with a 70° angle to achieve total internal reflection. Laser lines were selected with interference filters (Semrock) in a motorized filter wheel equipped with a shutter (Sutter Instruments) blocking the beam between image captures. Emission light was detected at 530/50 nm (Semrock interference filters) for GFP or 597LP (Melles Griot glass filter) for mCherry using a CCD camera (Hamamatsu ORCA-AG) controlled by MetaFluor software (Molecular Devices). For high resolution TIRF imaging we used an Eclipse TiE microscope (Nikon) equipped with a TIRF illuminator and a 60×/1.45-NA or 100×/1.49-NA objectives as previously described (Xie et al., 2019). Confocal microscopy was performed on an Eclipse TE2000 microscope (Nikon) equipped with a Yokogawa CSU-10 spinning disc confocal unit and a 100×/1.49-NA plan Apochromat objective (Nikon) (Idevall-Hagren et al., 2013). Briefly GFP and mCherry were excited by 491-nm and 561-nm DPSS lasers (Cobolt, Sweden) and were detected through 530/50 nm interference filter and 597LP filter, respectively, through a black-illuminated EM-CCD camera (DU-888; Andor Technology).

### Fluorescence recovery after photobleaching (FRAP)

TIRF-FRAP was performed on a Nikon TiE microscope equipped with an iLAS2 TIRF illuminator for multi-angle patterned illumination (Cairn Research) and a 100X1.49-NA Apo-TIRF objective. Excitation light for GFP and mCherry was delivered by 488-nm and 561-nm diode-pumped solid-state lasers with built-in acousto-optical modulators, and light for bleaching was delivered by a 405-nm DPSS laser (all from Coherent). Fluorescence was detected with a back-illuminated EMCCD camera (DU-897, Andor Technology) controlled by MetaMorph (Molecular Devices). Emission wavelengths were selected with filters (527/27 nm for GFP and 590 nm long-pass for mCherry) mounted in a filter wheel (Sutter Instruments). Cells were mounted in an open perfusion chamber with temperature held at 37°C. Following a 10 s baseline acquisition, a 3×3 μm area within the cells was exposed to a 100 ms 405-nm light pulse to bleach the fluorophores, followed by continued acquisition for 120 s at 4 fps. Calculations of the mobile fraction (*Mf*) of each fluorescent protein was performed using the following formula:

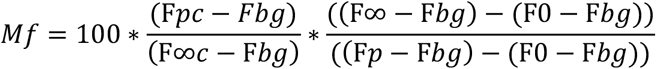

F*pc* (whole cell pre-bleach intensity), F*p* (bleach ROI pre-bleach intensity), F*∞c* (asymptote of fluorescence recovery of the whole cell), F*bg* (mean background intensity), F*∞* (asymptote of the bleach ROI), F*0* (bleach ROI post-bleach intensity).

### Western blot analysis

After 3-time wash with phosphate-buffered salin (PBS), MIN6 cells were homogenized and lysed in RIPA buffer (50mM Tris-HCl, pH=7.4, 1% NP-40, 0.5% Na-deoxycholate, 0.1% SDS, 150mM EDTA), placed on ice for 30 min or agitated on a rotator at 4°C for 30 min. Cell lysates were mixed with 3×sample buffer (6% SDS, 15% 2-mercaptoethanol, 30% glycerol, 0.006% bromophenol blue, 0.15 M Tris-HCl) and boiled at 95°C for 10 min. Protein content was determined by a detergent-compatible protein assay (Bio-Rad, Hercules, CA, USA) and proteins were separated by SDS-PAGE (5-20%) and blotted onto PVDF membrane using semi-dry transfer. Membranes were blocked in 4% milk dissolved in TBS-Tween (0.1% Tween 20). The following antibodies and dilutions were used: anti-TMEM24 (Bethyl Laboratories,1:1000), anti-GAPDH (Cell Signaling Technology,1:1000), anti-Rabbit secondary antibody (GE healthcare,1:10000). Membranes were developed by the Odyssey Fc Imaging system (LI-COR Bioscience).

### Quantitative RT-PCR

mRNA from control and TMEM24 knockdown cells were extracted using the NucleoSpin RNAPlus kit (Macherey-Nagel). RT-PCR was performed by QuantiTect SYBR Green RT-PCR kit (Qiagen) using the following primer: GAPDH-fwd, ACTCCACTCACGGCAAATTC-3; GAPDH-rev,TCT CCATGGTGGTGAAGACA; TMEM24-fwd, CGCCCAGAACTCAGCCTAAA, TEME24-rev GGGTAGGTCTGGGGATGGAT. PCR reaction was done by Light Cycler 2.0 (Roche). Results are presented as ΔΔC_t_, normalized by GAPDH expression in control and TMEM24 KD cells.

### Insulin secretion measurements

Insulin secretion was measured from monolayers of MIN6 cells grown in 12-well plates. 4×10^5^ control or TMEM24 knockdown cells were prepared as describe above in 12-well plates 72 hours before secretion measurements. Cells were pre-incubated in 3 mM glucose imaging buffer at 37 ℃ for 30 min. Cells were subsequently incubated in buffers containing 3 mM glucose, 20 mM glucose or 3 mM glucose supplemented with 30 mM KCl, immediately followed by buffer collection. In the end, cells were released by the addition of 80 μl typsin and mixed with 120 μl complete growth medium. 100 ul of the collected cell suspension was mixed with 100 μl acidic ethanol, sonicated on ice and neutralized by the addition of 900 μL Tris buffer (pH 8) to determine insulin content. The insulin concentration in the samples was measured by a mouse insulin AlphaLISA kit (Perkin-Elmer).

### Oxygen consumption rates measurement

The oxygen consumption rate (OCR) in MIN6 cells were monitored by an Extracellular Flux Analyzer (XFe96, Agilent Tchnologies, CA, USA). 3×10^4^ cells (wildtype and TMEM24 KO) were seeded in each well of XFe96 microplates and cultured for 24 additional hours. Cells were thereafter pre-incubated with Seahorse XF DMEM (Agilent Technologies) containing 3 mM glucose (pH 7.4) for 1h at 37°C before the microplate was inserted into the XFe96 Analyzer. For each experiment, 4–6 replicates of each treatment were measured. OCR at 3 mM glucose was measured for 30 minutes, followed by another 30 minutes with either 3 mM or 20 mM glucose. Then, the proportions of respiration driving ATP synthesis and proton leak were determined by blocking ATP synthase with 2 μM oligomycin (Sigma-Aldrich). Subsequently, 2 μM of the mitochondrial uncoupler cyanide-p-trifluoromethoxyphenylhydrazone (FCCP) was added to determine the maximal respiratory capacity. Lastly, 2 μM rotenone and 5 μM antimycin A were added together to block transfer of electrons from the mitochondrial respiratory chain complex I and III to determine the remaining non-mitochondria-dependent respiration. To calculate the mitochondrial respiration, non-mitochondrial OCR was subtracted from the total OCR. Data was normalized to protein content which was determined by the DC protein assay (Bio-Rad Laboratories, USA).

### Measurements of mitochondrial membrane potential

The mitochondrial membrane potential (ΔΨ_m_) was monitored by the lipophilic cationic dye tetramethylrhodamine methyl ester (TMRM) via fluorescence time-lapse imaging using an inverted microscope (Eclipse TE2000U; Nikon, Kanagawa, Japan). The epifluorescence microscope is equipped with a high-power LED light source (Omicron LedHUB; Photonlines Ltd, Newcatle, UK) which, connected with a 5-mm diameter liquid light guide, provided excitation light at 540 nm. Emission was measured at 560 nm (5 nm half-bandwidth) and the fluorescence signal was detected by an Evolve EMCCD camera (Photometrics, Arizona, USA). MetaFluor software (Molecular Devices Corp.) allowed to control the microscope setup and to acquire images every 5 seconds. Prior to imaging, the cells were seeded onto 25 mm round coverslips and loaded with TMRM at 10 nM during a 30-min incubation at 37 °C in an imaging buffer containing 125 mM NaCl, 5 mM KCl, 1.3 mM CaCl_2_, 1.2 mM MgCl_2_, and 25 mM HEPES with pH adjusted to 7.40 with NaOH. The coverslips were placed on the bottom of an open Sykes-More chamber. On the top of the coverslip, a thin 25-mm diameter stainless steel plate with a 4-mm wide and 7-mm long opening pressed the 1-mm thick silicon rubber gasket with identical dimensions and central opening to the coverslip. The temperature of the chamber holder and the CFI S Fluor 40 × 1.3 numerical aperture oil immersion objective (Nikon) was stable at 37°C during the experiment using custom-built thermostats. Fixed on the stainless-steel plate, inlet and outlet cannulas maintained a laminar superfusion at a rate of 2.0 ml/min with the imaging buffer containing 10 nM TMRM.

### Measurement of cytoplasmic, mitochondrial and endoplasmic reticulum [Ca^2+^]

Cytosolic [Ca^2+^] was measured on an epifluorescence microscope setup (described above) using the ratiometric dye Fura-2 or the green-fluorescent dye Cal520. The cells were preincubated for 30 min at 37°C in imaging buffer supplemented with 1 μM of the indicator, followed by repeated washing in imaging buffer and imaging. Mitochondrial Ca^2+^ was measured using the genetically-encoded red fluorescent, low affinity indicator mito-LAR-GECO (Wu et al., 2014), while ER Ca^2+^ was measured using the FRET-based indicator D4ER (Ravier et al., 2011). Both indicators were delivered to cells by transient transfection as described above, and experiments were performed 18-36h post-transfection.

### Morphometric analysis of mitochondria

To determine the morphology of the mitochondria, wildtype MIN6 cells and TMEM24 KO cells expressing mApple-Tomm20 were observed under the spinning-disc confocal microscope. The images were analyzed with the open-source image analysis software CellProfiler (Carpenter et al., 2006) using several modules that were placed in a sequential order to create a flexible image analysis pipeline. First, processing filters were applied to enhance the fluorescence signal of the regions that display higher intensity relative to its immediate neighborhood. This allowed the better identification of the mitochondria as separate objects during the following module. The Otsu’s automatic threshold method permitted the assignment of the threshold value by including the pixels of the image either in the “background” class or the “foreground” class. The objects/mitochondria detected using the pipeline were not de-clumped to enable the identification of potential network formation, and the size and shape of the mitochondria was determined and the data was exported to Microsoft Excel. Eccentricity, which distinguish mitochondria based on shape from tubular (1) to circular (0), was used as an overall determinant of the mitochondria shape.

### Image analysis

TIRF microscopy and confocal microscopy images were analyzed offline by Fiji (Schindelin et al., 2012). To determine fluorescence changes, the regions of interest and background regions were first manually identified. Fluorescence intensity changes within these regions were recorded and the data was exported to Excel. All data points were background corrected and normalized to the initial fluorescence intensity (F/F_0_).

### Statistical analysis

One-way ANOVA followed by Tukey’s Post hoc test, Mann-Whitney U-test (for non-parametric data) or Student t-test were used.

## RESULTS

### TMEM24 plasma membrane binding is controlled by DAG and Ca^2+^

To better understand the role of TMEM24 in the regulation of β-cell function, we investigated conditions that promote TMEM24 dissociation from the plasma membrane. Elevation of the glucose concentration from 3 to 20 mM in clonal MIN6 β-cells resulted in regular cytosolic Ca^2+^ oscillations with elevations mirrored by synchronized TMEM24-GFP dissociations, seen as reductions in plasma membrane-proximal fluorescence by TIRF microscopy (Fig. 1A-C). Direct depolarization with 30 mM KCl also caused dissociation of TMEM24 from the plasma membrane, consistent with voltage-dependent Ca^2+^ influx being the trigger for dissociation (Fig. 1B-D). Similar TMEM24 dissociation was also seen in response to carbachol, which activate phospholipase C and trigger IP3-mediated release of Ca^2+^ from the ER (Fig. 1B, C). Passive depletion of ER Ca^2+^ using the SERCA inhibitor cyclopiazonic acid (CPA) also triggered TMEM24 dissociation, although less prominent than in response to carbachol (Fig. 1B, C). TMEM24 dissociation has previously been shown to depend on PKC-mediated phosphorylation of C-terminal residues in the molecule (Lees et al., 2017). Although Ca^2+^ is a potent activator of PKC, some isoforms also require diacylglycerol (DAG). To determine to what extent DAG is involved in the spatial control of TMEM24, we stimulated MIN6 cells with 1 μM DAG analogue Phorbol 12-Myristate 13-Acetat (PMA). This resulted in an immediate dissociation of TMEM24 from the plasma membrane without apparent change in the cytosolic Ca^2+^ concentration (Fig. 1B, C and Suppl. Fig. 1A). To further uncouple DAG formation from Ca^2+^ changes we first exposed cells to CPA to deplete ER Ca^2+^ stores, followed by addition of carbachol to activate PLC and increase plasma membrane DAG. CPA caused an increase in cytosolic Ca^2+^ and a slight dissociation of GFP-TMEM24 from the plasma membrane. The subsequent addition of carbachol resulted in a pronounced dissociation of TMEM24 that occurred in the absence of noticeable changes in cytosolic Ca^2+^ concentration (Suppl. Fig. 1B). These results indicate that TMEM24 is spatially controlled by both Ca^2+^ and DAG. We also noticed that whereas carbachol caused a homogenous dissociation of TMEM24 from the plasma membrane (Fig. 1E, right panels), the dissociation in response to direct depolarization with KCl were instead incomplete, more heterogenous and differed between sub-regions of the plasma membrane within the same cell (Fig. 1E, left panels). This pattern resembles that of a diacylglycerol (DAG) biosensor (GFP-C1aC1b_PKC_) reporting DAG formation after autocrine activation of purinergic P2Y1 receptors by ATP co-secreted with insulin from the cells (Wuttke et al., 2013) (Suppl. Fig. 1C, D). Because of the close interplay between DAG and Ca^2+^, we determined the effect of known Ca^2+^ concentrations on TMEM24-mCherry plasma membrane binding in α-toxin-permeabilized MIN6 cells in which changes in Ca^2+^ could easily be uncoupled from those in DAG. We found that dissociation of TMEM24 occurred with an EC_50_ 358±70 nM (n=13) and maximal effect at 30 μM (Fig. 1F, G). Addition of 1 μM PMA caused pronounced dissociation of TMEM24 from the plasma membrane and subsequent stepwise increase in the Ca^2+^ concentration had little effect on TMEM24 plasma membrane binding (Suppl. Fig. 1E). For comparison, the plasma membrane binding of E-Syt1-mCherry, another Ca^2+^-dependent ER-PM contact protein, was triggered at 1 μM Ca^2+^ and maximal at 10 μM Ca^2+^ (EC_50_ 1.99±0.51 μM, n=6) (Fig. 1G). When comparing the plasma membrane fluorescence intensity of TMEM24-mCherry in intact cells and following permeabilization in a Ca^2+^-free buffer, we found that removal of Ca^2+^ promoted further association of TMEM24 with the plasma membrane (13±5% increase in plasma membrane TMEM24-mCherry fluorescence, n=22) (Fig. 1H) while the opposite was observed for GFP-E-Syt1 (29±4% drop in plasma membrane fluorescence, n=31). These results indicate that TMEM24 plasma membrane binding may be partially reduced already at resting Ca^2+^ concentrations. Fluorescence recovery after photobleaching (FRAP) analysis revealed that TMEM24-GFP, although enriched at ER-PM contacts, exhibited pronounced dynamics with a large proportion belonging to a mobile fraction (Fig. 1I). Taken together, these results show that the association of TMEM24 with the plasma membrane is regulated by both DAG and Ca^2+^, and that dissociation is triggered already by modest nanomolar Ca^2+^ concentrations, resulting in relatively weak interactions between TMEM24 and the plasma membrane.

**Figure 1.**
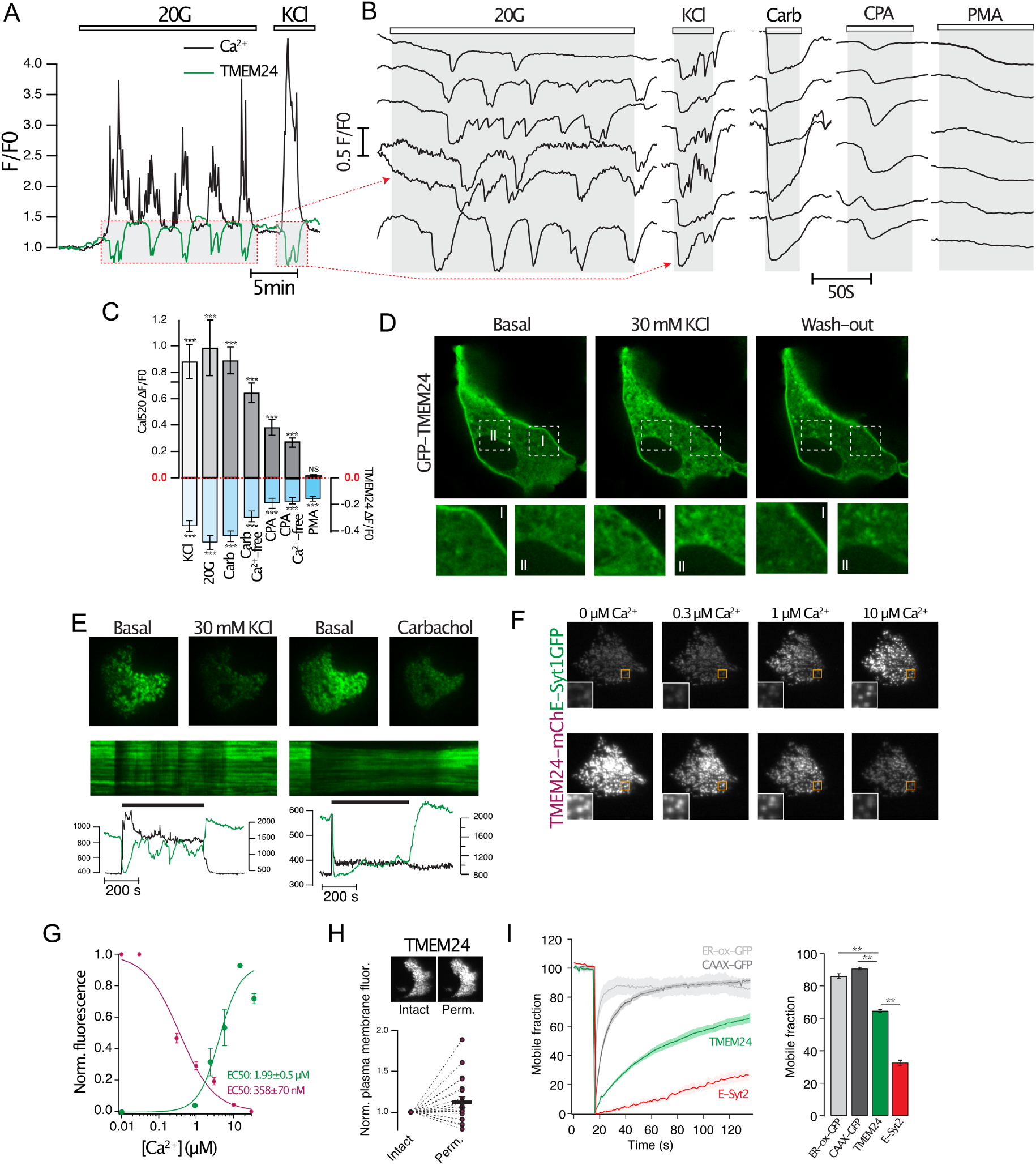
TMEM24 plasma membrane binding is controlled by DAG and Ca^2+^. (A) TIRF microscopy recording of Ca^2+^ indicator Cal520 (black) and TMEM24-EGFP (green) fluorescence from a single MIN6 cell stimulated with 20 mM glucose (20G) and 30 mM KCl. (B) Representative TIRF microscopy recordings from TMEM24-EGFP expressing MIN6 cells in response to 20 mM glucose, 30 mM KCl, 100 μM carbachol, 100 μM CPA and 1 μM PMA. (C) Quantification of Ca^2+^ increases (grey) and the corresponding magnitude of TMEM24-EGFP dissociation from the plasma membrane (blue), n=20 cells for each group. *** P<0.001 for comparison to 0. (D) Confocal microscopy images of a MIN6 cell expressing TMEM24-EGFP. Images were taken before, during and 30 s after depolarization. Content within the dashed boxes are magnified below. (E) TIRF microscopy images of two MIN6 cells expressing TMEM24-EGFP. Pictures to the left shows a cell before and 30 s after the addition of 30 mM KCl and pictures to the right shows a cell before and 30 s after addition of 100 μM carbachol. Shown below are kymographs of TMEM24-EGFP fluorescence from lines drawn across each cell and TIRF microscopy recordings from the cells (black trace, Cal520 fluorescence; green trace, TMEM24-EGFP fluorescence) (F) TIRF microscopy images of MIN6 cells loaded with the Ca^2+^ indicator Cal520 and expressing E-Syt1-GFP (top) or TMEM24-mCherry (bottom) following a-toxin permeabilization and exposure to the indicated Ca^2+^ buffers. (G) Dose-response curves of Ca^2+^ induced TMEM24-mCherry (magenta) and E-Syt1-GFP (green) fluorescens changes in α-toxin permeabilized MIN6 cells (n=42 cells). (H) TIRF microscopy images of a MIN6 cell expressing TMEM24-mCherry before (intact) and after atoxin permeabilization (perm). Quantifications below show the plasma membrane fluorescence change that occurred following a-toxin permeabilization in a Ca^2+^-deficient buffer. (I) TIRF microscopy recordings of ER-oxGFP (ER luminal protein; light grey), GFP-CaaX (prenylated protein anchored in the plasma membrane; dark grey), TMEM24-EGFP (green) and E-Syt2-GFP (red) fluorescence recovery after photobleaching (FRAP). Bar graph to the right shows the mobile fraction of each fluorescently-tagged protein. All data are presented as mean±SEM, ***p<0.001

### TMEM24 is not required for glucose-stimulated insulin secretion

TMEM24 has previously been reported to be required for glucose-stimulated Ca^2+^ influx and insulin release, although the mechanisms are unclear (Lees et al., 2017). To examine its role in more detail, we reduced TMEM24 expression in MIN6 cells by siRNA-mediated transient knockdown. such knockdown resulted in 61.2±6.3% (n=4, P<0.01) reduction in TMEM24 expression as assessed by Western blotting (86.7±5.2% reduction as assessed by qRT-PCR) (Fig. 2A). We next determined the impact of reduced TMEM24 expression on Ca^2+^ handling using the fluorescent Ca^2+^ indicator Cal520. In control cells, elevation of the glucose concentration from 3 to 20 mM resulted in rapid lowering of the cytosolic Ca^2+^ concentration (−17.1±0.9%, n=148) due to ATP-driven sequestration of the ion into the ER (ref), followed after 2.8±0.09 minutes by a pronounced Ca^2+^ increase and sustained elevation (Fig. 2B, C, E). Most cells showed slow, regular Ca^2+^ oscillations in the continued presence of 20 mM glucose, while a smaller number showed more irregular responses (Fig. 2B). In TMEM24 KD cells, the initial Ca^2+^ lowering in response to 20 mM glucose was less pronounced (−12.4±0.8%, n=137, P=0.0016 for comparison to control) and was followed after 3.31±0.11min (n=141, P=0.003 for comparison to control) by a sustained rise of Ca^2+^ with regular oscillations (Fig. 2B, C, E). The time-average increase in Ca^2+^ was slightly higher in TMEM24 KD cells compared to controls (Fig. 2C, D). Overexpression of TMEM24 resulted in an exaggerated initial lowering of the cytosolic Ca^2+^ concentration in response to glucose and a slightly increased sustained Ca^2+^ response when compared to non-transfected control cells (Fig. 2C-E). To ascertain that the relatively mild effect on Ca^2+^ handling in TMEM24 KD cells was not due to remaining low TMEM24 expression, we generated TMEM24 knockout MIN6 cell lines using CRISPR/Cas9 (Fig. 2F). Both control and TMEM24 KO cells exhibited subsequent oscillations in the cytosolic Ca^2+^ concentration, although the time-averaged Ca^2+^ increase was reduced by 21±2% (n=497, P=1.47E-9) in the TMEM24 KO cells (Fig. 2G-I). Similar to TMEM24 KD, the KO cell line also exhibited strongly suppressed initial Ca^2+^ lowering in response to glucose (Fig. 2J, K). Both TMEM24 KD and TMEM24 KO cells exhibited Ca^2+^ responses to glucose that contained a component of rapid Ca^2+^ transients superimposed on top of the regular, slow Ca^2+^ oscillations (Fig. 2B, G).

**Figure 2.**
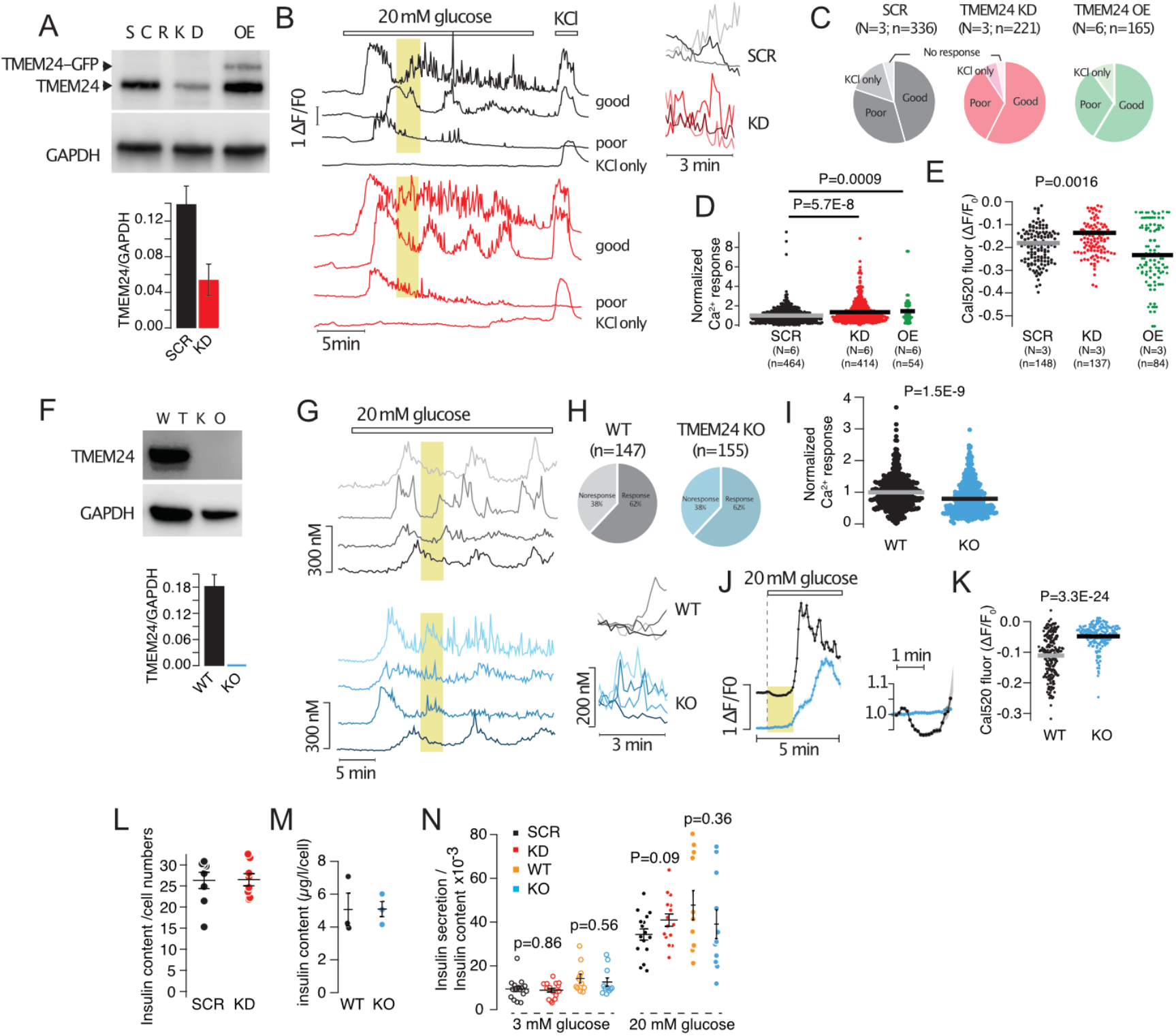
TMEM24 is not essential for glucose-induced Ca^2+^ signaling or insulin secretion. (A) Western blot of lysates from MIN6 cells transfected with control (SCR) or anti-TMEM24 siRNA (KD) alone or together with TMEM24-EGFP (+TMEM24-EGFP) probed with anti-TMEM24 and GAPDH antibodies. Quantifications of densitometric measurements are shown below (n=4). (B) Representative Ca^2+^ recordings (Fluo-4) from control (black) and TMEM24 KD (red) MIN6 cells in response to 20 mM glucose and 30 mM KCl. The cells were divided into three groups based on the type of response: ‘good’; continuous response to glucose and robust response to KCl, ‘poor’; initial, but not sustained, response to glucose and robust response to KCl, ‘KCl only’; no response to glucose but robust response to KCl. The boxed yellow areas are shown to the right on an expanded time-scale. (C) Pie chart showing the distribution of Ca^2+^ responses in control (black), TMEM24 KD (red) and TMEM24-EGFP expressing (green) MIN6 cells. (D) Time-average Fluo-4 fluorescence change in response to 20 mM glucose in control (SCR, black), TMEM24 KD (KD, red) and TMEM24-EGFP expressing (OE, green) MIN6 cells. (E) Quantifications of the initial, glucose-induced lowering of Fluo-4 fluorescence in control (SCR, black), TMEM24 KD (KD, red) and TMEM24-EGFP expressing (OE, green) MIN6 cells. (F) Western blot of lysates from wildtype and TMEM24 KO MIN6 cells probed with anti-TMEM24 and GAPDH antibodies. Quantifications of densitometric measurements are shown below (n=2). (G) Representative Ca^2+^ recordings (Fura-2) from control (black) and TMEM24 KO (blue) MIN6 cells in response to 20 mM glucose. The cells were divided into two groups based on the absence or presence of Ca2+ increase in response to glucose. The boxed yellow areas are shown to the right on an expanded time-scale. (H) Pie chart showing the distribution of Ca^2+^ responses in wildtype (black), TMEM24 KO (blue) MIN6 cells. (I) Time-average Ca^2+^ response to 20 mM glucose in wildtype (black) and TMEM24 KO (blue) cells. (J) Initial Ca^2+^ response to 20 mM glucose in wildtype (black) and TMEM24 KO (blue) cells. Data presented as means±SEM for 63 (wildtype) and 52 (KO) cells from one experiment. The yellow area is shown to the right on an expanded time-scale. (K) Quantifications of the initial, glucose-induced lowering of Cal520 fluorescence in wildtype (black) and TMEM24 KO (blue) MIN6 cells. (L,M) Insulin content in control and TMEM24 KD cells (L) and in wildtype and TMEM24 KO cells (M). (N) Insulin secretion in control (black), TMEM24 KD (red), wildtype (yellow) and TMEM24 KO cells in response to 3 mM and 20 mM glucose.

Given the relatively small of effect of TMEM24 KD or KO on glucose-induced Ca^2+^ influx observed here, we decided to reinvestigate the previously proposed fundamental role of TMEM24 in insulin secretion (Lees et al., 2017; Pottekat et al., 2013). Using insulin ELISA we found that basal secretion at 3 mM glucose and that stimulated by 20 mM were similar in control and TMEM24 KD and KO cells (Fig. 2N). There was also no difference in insulin content between control cells and TMEM24 KD or KO cells (Fig. 2L, M). These results indicate that TMEM24 is dispensable for the acute regulation of glucose-stimulated insulin secretion.

### TMEM24 controls ER Ca^2+^ homeostasis

Since we observed a slightly impaired Ca^2+^ response to glucose in TMEM24 KO cells, we tested to what extent Ca^2+^ influx in response to direct depolarization was affected by the loss of TMEM24. The resting cytosolic Ca^2+^ concentration was similar in wildtype and TMEM24 KO cells. Depolarization with 30 mM KCl resulted in an immediate rise of cytosolic Ca^2+^ that averaged 361±5 nM (n=298) in wildtype cells. This response was increased to 441±8 nM (n=317, P=3.12E-16) in TMEM24 KO cells (Fig. 3A, B). Consistent with more Ca^2+^ entering the cells in response to depolarization, we also found that insulin secretion in response to acute depolarization was increased by 60% in TMEM24 KO cells (n=12, P=0.019) (Fig. 3G). Similar results were obtained from cells where TMEM24 expression had instead been reduced by siRNA (Fig. 3D, E, H). Several observations made in TMEM24 KO cells, such as lack of initial Ca^2+^-lowering effect of glucose and the appearance of irregular Ca^2+^ spikes, pointed to potential changes in the ability of the ER to sequester and mobilize Ca^2+^. To test this more directly, we mobilized Ca^2+^ from the ER by the SERCA inhibitor CPA in the absence of extracellular Ca^2+^ while measuring changes in the cytosolic Ca^2+^ concentration. The addition of CPA resulted in a 32±1 nM (n=298) increase in the cytosolic Ca^2+^ concentration, reflecting the release of Ca^2+^ from the ER. This release was increased to 48±2 nM (n=306, P=9.8E-12) in TMEM24 KO cells (Fig. 3I, J). The addition of 10 mM Ca^2+^ to the extracellular buffer triggered store-operated Ca^2+^ entry, which was not different in the two cell lines (Fig. 3K). Similar results were obtained when ER Ca^2+^-store depletion was instead triggered by the addition of thapsigargin in the presence of extracellular Ca^2+^, and the response in TMEM24 KO cells was normalized by the re-expression of wildtype TMEM24 (Fig. 3L). Direct measurements of ER Ca^2+^ using the FRET-based Ca^2+^ sensor D4ER confirmed that the ER of TMEM24 KO cells contained more Ca^2+^ than that of wildtype cells under resting conditions, and also that more Ca^2+^ was released from the ER following SERCA inhibition with CPA (Fig. 3M, N). We speculated that perhaps the larger rise of cytosolic Ca^2+^ in response to depolarization in TMEM24 KO cells might be due to simultaneous Ca^2+^-induced Ca^2+^-release from the ER. To test this alternative, we performed experiments where cytosolic Ca^2+^ was measured following two brief applications of 30 mM KCl, where the second application was preceded by SERCA inhibition with thapsigargin. Consistent with previous observations (Chen et al., 2003) prevention of Ca^2+^ sequestration into the ER by SERCA inhibition resulted in a more pronounced depolarization-induced Ca^2+^ increase in wildtype cells (Fig. 3N, O). This augmentation was even more apparent in TMEM24 KO cells, and could be restored to the level of wildtype cells by the re-expression of TMEM24 (Fig. 3P, Q). These results indicate that Ca^2+^ sequestration into the ER may be a way to compensate for excess Ca^2+^ increase in TMEM24 KO cells rather than being the cause of it.

**Figure 3.**
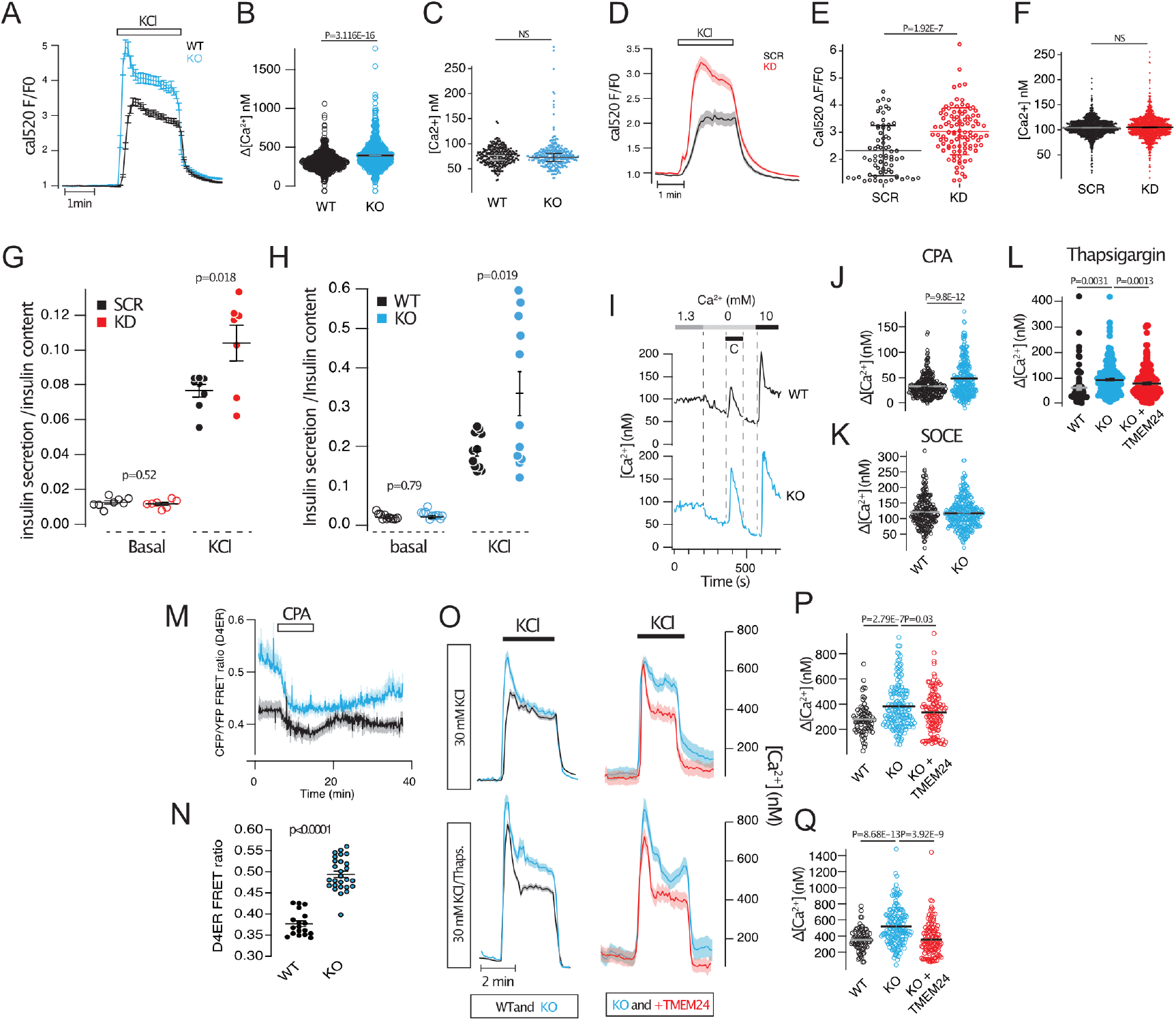
TMEM24 regulate ER Ca^2+^ homeostasis. (A) Depolarization-induced Ca^2+^-influx in wildtype (black) and TMEM24 KO (blue) cells. Traces show mean±SEM for 41 (SCR) and 54 (KD) cells from one experiment. (B) Average cytosolic Ca^2+^ concentration change in response to 30 mM KCl in wildtype (black, n=298) and TMEM24 KO (blue, n=397) cells. (C) Resting Ca^2+^ concentration in wildtype (black, n=512) and TMEM24 KO (blue, n=534) cells. (D) Depolarization-induced Ca^2+^-influx in control (SCR, black) and TMEM24 KD (KD, red) cells. Traces show mean±SEM for 22 (SCR) and 19 (KD) cells from one experiment. (E) Average Cal520 fluorescence increase in response to 30 mM KCl in control (black, n=73) and TMEM24 KD (red, n=101) cells. (F) Resting Ca^2+^ concentration in control (black) and TMEM24 KD (red) cells. (G) Insulin secretion in control (black) and TMEM24 KD (red) cells exposed to buffers containing basal (5.5 mM) and depolarizing (30 mM) concentrations of KCl (n=8). (H) Insulin secretion in wildtype (black) and TMEM24 KO (blue) cells exposed to buffers containing basal (5.5 mM) and depolarizing (30 mM) concentrations of KCl (n=12). (I) Representative Fura-2 recordings showing cytosolic Ca2+ concentration changes in wildtype (black) and TMEM24 KO (blue) cells following ER-store depletion with CPA and SOCE upon re-addition of extracellular Ca2+. (J) Quantifications of the increase in cytosolic Ca2+ in wildtype (black, n=296) and TMEM24 KO cells (blue, n=306) following addition of CPA shows that TMEM24 KO cells release more Ca2+ from the ER. (K) Quantifications of the increase in cytosolic Ca2+ in wildtype (black, n=296) and TMEM24 KO cells (blue, n=306) following SOCE shows that there is no difference between wildtype and TMEM24 KO cells. (L) Quantifications of the increase in cytosolic Ca2+ in response to thapsigargin shows that more Ca2+ is released from the ER in TMEM24 KO cells (blue, n=270) compared to wildtype cells (black, n=77), and that this can be rescued by the re-expression of TMEM24 (red, n=248). (M) Measurements of ER Ca2+ using D4ER shows that the resting ER Ca2+ concentration is higher in TMEM24 KO cells (blue) compared to wildtype cells (black). (N) Resting D4ER FRET ratios in WT (black; n=18) and TMEM24 KO (blue; n=27) cells (P<0.0001). (O) Average cytosolic Ca2+ concentration changes in wildtype (black), TMEM24 KO (blue) and TMEM24 KO with re-expression of TMEM24 (red) in response to 30 mM KCl alone (top row) or after thapsigargin addition (bottom row). Data are averages from 32-43 cells from 1 experiment per condition. (P) Quantifications of the increase in cytosolic Ca2+ in wildtype (black, n=77) and TMEM24 KO cells (blue, n=277) following depolarization shows that there is a larger increase in TMEM24 KO cells that can be normalized by the re-expression of TMEM24 (red, n=270). (Q) Quantifications of the increase in cytosolic Ca2+ in wildtype (black, n=77) and TMEM24 KO cells (blue, n=277) following depolarization in the presence of thapsigargin shows that there is a larger increase in TMEM24 KO cells that can be normalized by the re-expression of TMEM24 (red, n=270).

### TMEM24 regulates mitochondrial Ca^2+^ handling and ATP production

Because Ca^2+^ influx, extrusion and organellar sequestration all depends on ATP generated primarily via mitochondrial metabolism, we decided to complement the Ca^2+^ imaging with direct measurement of glucose metabolism using the Seahorse XF technique. As expected from the Ca^2+^ imaging data, TMEM24 knockout had little effect on the resting oxygen consumption rate (OCR) (Fig. 4A). In contrast, the accelerated OCR induced by a rise of the extracellular glucose concentration from 3 to 20 mM was markedly impaired in TMEM24 KO cells (Fig. 4B). However, neither the proton leak nor the maximal OCR was different between wildtype and TMEM24 KO cells, indicating that there was no gross impairment in overall mitochondrial function (Fig. 4C-F). This conclusion was supported by lack of apparent changes in mitochondrial morphology in TMEM24 KO cells as assessed by confocal microscopy of cells expressing mitochondria-targeted mApple (Tomm20-mApple) (Fig. 4G, H). Next, we measured the mitochondrial membrane potential in wildtype and TMEM24 KO cells using the fluorescent membrane potential indicator TMRM. Application of the uncoupler FCCP resulted in the immediate loss of TMRM fluorescence, which reflects depolarization of the inner mitochondrial membrane. This response was significantly smaller in TMEM24 KO cells, indicating that these cells already have partially depolarized mitochondria (Fig. 4I, J). The mitochondrial membrane potential does not only control ATP production by directly affecting the electron transport chain but also by regulating the amounts of Ca^2+^ that are taken up and extruded by the organelle. To determine whether TMEM24 might be involved in the regulation of mitochondrial Ca^2+^ we measured concentration changes in response to both CPA-mediated release of Ca^2+^ from the ER and depolarization-induced Ca^2+^ influx in wildtype and TMEM24 KO cell using mitochondrially targeted LAR-GECO1.2. This low-affinity sensor has a K_d_ for Ca^2+^-binding of 12 μM, and its fluorescence is expected to increase little under normal conditions, since depolarization-induced Ca^2+^ influx typically results in Ca^2+^ concentrations in the low μM range (see e.g. Fig. 3B). Consistently, KCl depolarization of wildtype cells triggered a robust increase in the cytosolic Ca^2+^ concentration, measured with the organic dye Cal520, but had little impact on mito-LAR-GECO1.2 fluorescence in the same cells (Fig. 4K, L). In contrast, depolarization caused a pronounced increase in mito-LAR-GECO1.2 fluorescence in TMEM24 KO cells, which was reduced to the level of wildtype cells by the re-expression of TMEM24 (Fig. 4K, L). Similar results were obtained when the Ca^2+^ increase was instead triggered by passive depletion from the ER through CPA-mediated SERCA inhibition (Fig. 4M, N). A possible explanation for these observations is that the resting mitochondrial Ca^2+^ concentration is higher in TMEM24 KO cells, thus bringing the concentration into a range better suited for detection with the low affinity sensor. Consistent with this hypothesis, we observed higher resting mito-LAR-GECO1.2 fluorescence in TMEM24 KO cells compared to wildtype cells (Fig. 4O).

**Figure 4.**
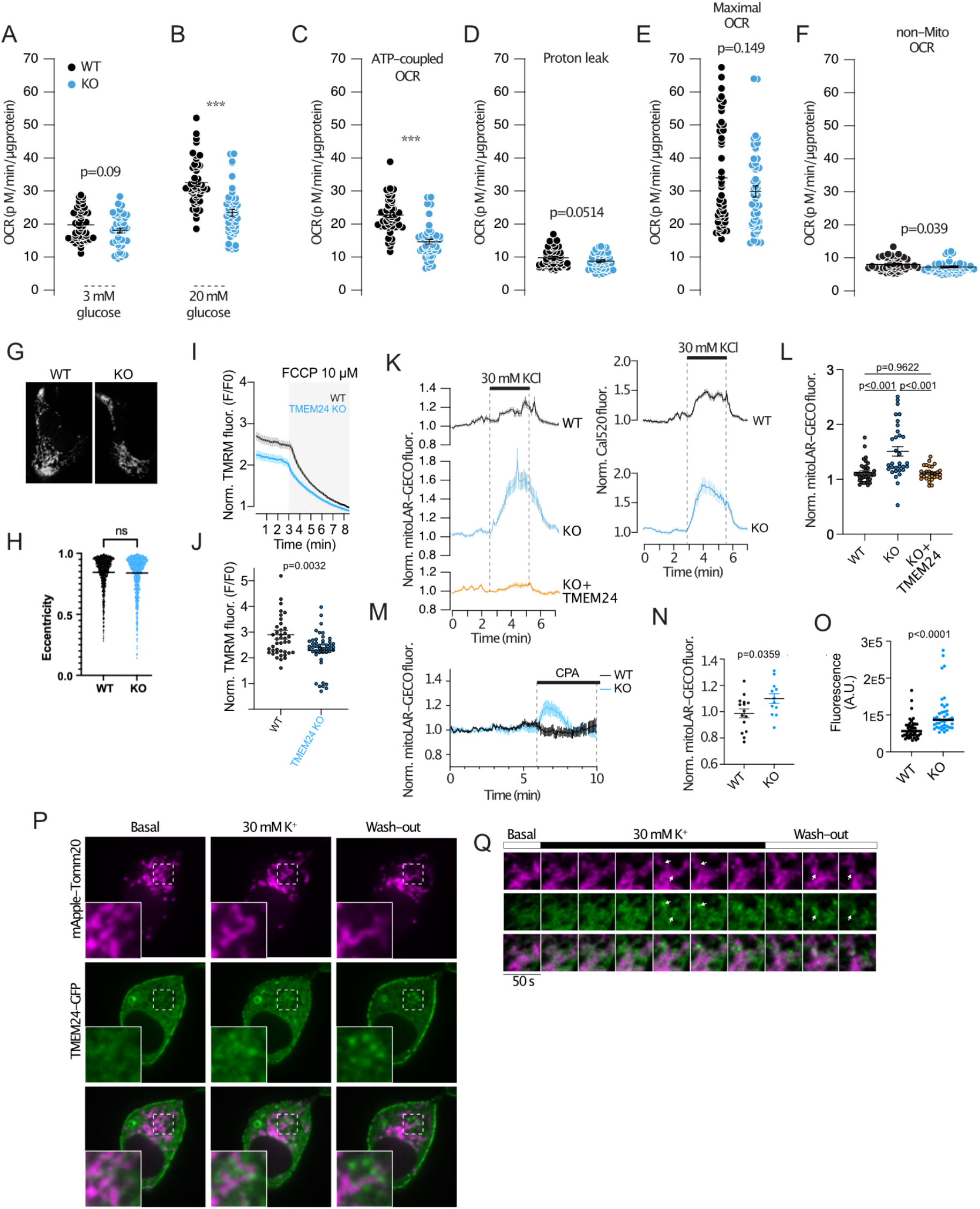
TMEM24 controls mitochondria Ca^2+^ handling and ATP production. (A) Oxygen consumption rate (OCR) in control (black) and TMEM24 KO (blue) cells at 3 mM glucose (n=40). (B) OCR in control (black) and TMEM24 KO (blue) cells at 20 mM glucose (n=40). OCR is reduced in TMEM24 KO cells (*** P<0.001). (C) ATP-coupled OCR in control (black) and TMEM24 KO (blue) cells (n=40) (*** P<0.001). (D) Proton leak in control (black) and TMEM24 KO (blue) cells (n=40). (E) Maximal OCR in control (black) and TMEM24 KO (blue) cells (n=40). (F) Non-mitochondrial OCR in control (black) and TMEM24 KO (blue) cells (n=40). (G) Confocal microscopy images of wildtype (left) and TMEM24 KO (right) cells expressing the mitochondrial marker Tom20-mApple. (H) Measurements of the eccentricity of identified mitochondria in wildtype (black) and TMEM24 KO (blue) cells (n=918 mitochondria from 50 cells for WT and n=1037 mitochondria from 55 cells for KO, P=0.3965). (I) Epifluorescence microscopy recordings of TMRM fluorescence from wildtype (black) and TMEM24 KO (blue) cells in response to 10 μM FCCP which causes dissipation of the inner mitochondrial membrane potential. Data presented are means ± SEM for 40 (WT) and 46 (KO) cells from 3 experiments. (J) Resting TMRM fluorescence values in wildtype and TMEM24 KO cells after normalization to the fluorescence intensity in the presence of 10 μM FCCP (n=40-46 cells; P=0.0032). (K) Epifluorescence microscopy recordings of mito-LAR-GECO (left column) and Cal520 (right column) fluorescence from wildtype cells (black; n=36 cells), TMEM24 KO cells (blue; n=32 cells) and TMEM24 KO cells with re-expression of TMEM24 (yellow; n=32 cells) (averages ± SEM; P<0.0001 for KO to WT and P=0.9622 for WT to rescue, One-way ANOVA). (L) Mito-LAR-GECO fluorescence change in response to 30 mM KCl in wildtype cells (black; n=40 cells), TMEM24 KO cells (blue; n=42 cells) and TMEM24 KO cells with re-expression of TMEM24 (yellow; n=29 cells). (M) Epifluorescence microscopy recordings of mito-LAR-GECO fluorescence from wildtype (black; n=15) and TMEM24 KO (blue; n=12) cells (averages ± SEM). (N) Mito-LAR-GECO fluorescence change in response to 100 μM CPA in wildtype (black) and TMEM24 KO (blue) cells (P=0.0359). (O) Resting mito-LAR-GECO fluorescence intensity in wildtype (black; n=50 cells) and TMEM24 KO (blue; n=43 cells) cells (P<0.0001). (P) Confocal microscopy images of a MIN6 cells expressing mApple Tomm20 (magenta) and TMEM24-GFP (green) under resting conditions, during stimulation with 30 mM KCl and following washout of the depolarizing stimuli. Boxed areas are magnified in the lower left corner and shows the appearance of TMEM24-GFP-positive puncta at mitochondria. (Q) Time-course of KCl-induced TMEM24 dissociation from the plasma membrane and association with mitochondria. Bars onto indicate addition and removal of KCl. Time interval between images is 50 s.

We next asked how an ER-localized protein that primarily engage in lipid transport at ER-PM contact sites could impact mitochondrial function. One possibility is that TMEM24 can function at other membrane contact sites. As we show here, a large part of the PM-bound pool of TMEM24 is dynamic even under resting conditions and could therefore participate in reactions at other cellular membranes, such as those of mitochondria. To test this alternative, we expressed GFP-tagged TMEM24 together with mitochondrially localized mApple-Tomm20 in wildtype MIN6 cells. Using confocal microscopy we did not observe any mitochondrial enrichment of TMEM24 under resting conditions, but clusters appeared at mitochondria when TMEM24 dissociation from the plasma membrane was stimulated by KCl-depolarization (Fig. 4P, Q). These clusters persisted and became even more striking when the bulk TMEM24 re-associated with the plasma membrane after terminating the depolarization (Fig. 4P, Q). These results show that TMEM24, in addition to acting at ER-PM contact sites, also may engage in reactions at ER-mitochondria contacts.

## DISCUSSION

Previous work has established the ER-localized lipid-transport protein TMEM24 as an important regulator of insulin secretion from pancreatic β-cells (Lees et al., 2017; Pottekat et al., 2013). Transient reduction of TMEM24 expression or knockout of the TMEM24 gene have been found to impair insulin secretion from clonal rat and mouse β-cells. The underlying mechanism has been proposed to involve reduced Ca^2+^-independent recruitment of insulin granules to the plasma membrane (Pottekat et al., 2013) or disturbed regulation of voltage-dependent Ca^2+^ influx through alterations in the plasma membrane lipid composition (Lees et al., 2017). In this work, we further explored the mechanisms underlying TMEM24-dependent regulation of insulin secretion. In contrast to previous studies, we found that reduced expression of TMEM24 has little impact on voltage-dependent Ca^2+^ influx and insulin secretion from β-cells. Instead, we identified TMEM24 as an important regulator of Ca^2+^ homeostasis at both the endoplasmic reticulum and the mitochondria, and show that this protein regulates mitochondrial ATP production.

TMEM24 is anchored to the endoplasmic reticulum through an N-terminal transmembrane domain and contains, in sequence, a lipid-binding SMP-domain, a C2-domain and a polybasic C-terminus. It is highly expressed in neuronal and endocrine tissue, including the pancreatic islets of Langerhans, where it is involved in tethering the ER to the plasma membrane (Lees et al., 2017; Sun et al., 2019). Its localization to the plasma membrane depends on electrostatic interactions between the C-terminus and cationic lipids in the plasma membrane, and neutralization of charged amino acids in TMEM24 by PKC-dependent phosphorylation results in TMEM24 dissociation. Using both intact and permeabilized cells, we found that the Ca^2+^-dependent dissociation of TMEM24 from the plasma membrane can be triggered already at nanomolar elevations of cytosolic Ca^2+^ by intracellular release or extracellular influx. Interestingly, we found that TMEM24 is already partially displaced from the plasma membrane under resting Ca^2+^ concentrations, perhaps due to constitutive PKC activity. This is consistent with findings from neuron-like cells where endogenous TMEM24 was found both at the plasma membrane and in the bulk ER under resting conditions (Sun et al., 2019). Such observations would support the hypothesis that TMEM24 acts at additional cellular locations, perhaps driven by interactions with acidic lipids in the membranes of organelles known to form contacts with the ER (Cohen et al., 2018; Eden et al., 2010). By locally photobleaching a small region of the plasma membrane in cells expressing TMEM24-GFP, we could also estimate the mobility of TMEM24 at ER-PM junctions. We found that TMEM24 weakly interacts with the plasma membrane, and that a large fraction was highly dynamic under resting conditions, which is in sharp contrast to Extended-Synaptotagmin-2, another SMP-domain-containing protein constitutively localized to ER-PM contact sites (Xie et al., 2019). This observation is also consistent with TMEM24 executing functions at other cellular locations than the plasma membrane.

In contrast to previous studies (Lees et al., 2017; Pottekat et al., 2013), we did not find support for a requirement of TMEM24 for normal insulin secretion. Both transient knockdown of TMEM24 using siRNA and CRISPR/Cas9-mediated knockout of TMEM24 had little effect on either glucose- or depolarization-induced Ca^2+^ influx and insulin secretion. If anything, cells with reduced expression performed slightly better than control cells. One previous study, in which TMEM24 expression was stably reduced by shRNA, showed impaired glucose-stimulated insulin secretion from both clonal rat INS1 and mouse MIN6 β-cells. However, the Ca^2+^ responses in these cells were normal, and secretion in response to direct depolarization was also unaffected by TMEM24 knockdown (Pottekat et al., 2013). CRISPR/Cas9-mediated knockout of TMEM24 in INS1 cells resulted in complete inhibition of both glucose-induced Ca^2+^ increases and insulin secretion, which was restored by re-expression of full-length TMEM24 (Lees et al., 2017). Although INS1 cells secrete insulin in response to glucose, the mechanism is likely different from that of primary β-cells in that it not only depends on K_ATP_-channel closure (Herbst et al., 2002) and voltage-dependent Ca^2+^ influx (Dorff et al., 2002). Insulin secretion may instead be triggered by Ca^2+^ released from intracellular stores, since addition of the ER Ca^2+^ ATPase (SERCA) inhibitor thapsigargin causes Ca^2+^ oscillations in these cells (Herbst et al., 2002). We found that TMEM24 knockout cells have increased Ca^2+^ accumulation in the ER, observed by both measurements of cytosolic Ca^2+^ following ER-store depletion and by direct measurements of ER Ca^2+^. It is not clear how TMEM24 contributes to ER Ca^2+^ homeostasis. The major route of Ca^2+^ uptake into the ER of β-cells is the SERCA (Roe et al., 1994) but we did not find any evidence for increased SERCA activity in TMEM24 KO cells. If anything, the activity was slightly reduced, as indicated by the lack of an initial Ca^2+^ lowering effect of glucose in TMEM24 KO cells, although this might also reflect the impaired ATP production or the steeper concentration gradient in these cells. There is a continuous leakage of Ca^2+^ from the ER, which results in rapid loss of Ca^2+^ upon SERCA inhibition, but the mechanism behind this leak is not clear. Numerous mediators of ER Ca^2+^ leak have been identified, including presenilin-1/2 and TMCO1 (Tu et al., 2006; Q. C. Wang et al., 2016). Reduced ER Ca^2+^ leak could explain the increased accumulation of Ca^2+^ in the ER of TMEM24 KO cells. Interestingly, loss of both presenilin-1 and TMCO1, like TMEM24, results in ER Ca^2+^ overload and in impaired mitochondria function (Tu et al., 2006; Q. C. Wang et al., 2016; X. Wang et al., 2019). Although TMEM24 unlikely functions as a Ca^2+^ channel, it may modulate other release mechanisms either through direct interactions or modulation of the lipid environment. Because of its dynamic nature, TMEM24 may provide means to acutely regulate ER Ca^2+^ permeability in response to increases in cytosolic Ca^2+^ or plasma membrane DAG concentrations.

In addition to its effect on ER Ca^2+^, we found that TMEM24 is involved in the regulation of mitochondria function. Increases in the cytosolic Ca^2+^ concentration causes the dissociation of TMEM24 from the plasma membrane and is followed by accumulation of TMEM24 at mitochondria. This is similar to other ER-localized lipid transport proteins, such as ORP5/8 and Vps13A/C, which has been shown to bind more than one organelle membrane (Galmes et al., 2016; Kumar et al., 2018). We also showed that the basal mitochondrial Ca^2+^ concentration is elevated in TMEM24 KO cells. However, these mitochondria are still able take up and release Ca^2+^ in response to changes in cytosolic Ca^2+^, arguing against defects in the major uptake and extrusion pathways. Consistent with the accumulation of Ca^2+^, we also found that the inner mitochondrial membrane was depolarized compared to mitochondria of wildtype cells and that the glucose-induced ATP production was impaired. It is currently not clear if it is the change in mitochondrial membrane potential that facilitates Ca^2+^ uptake via the mitochondrial calcium uniporter (MCU) complex or if the Ca^2+^ overload instead drives the depolarization. Irrespective of cause, it is likely that TMEM24 exert its effect on mitochondria at ER-mitochondria contact sites. These sites for Ca^2+^ and lipid exchange are important for controlling mitochondria Ca^2+^ levels, ATP production and mitochondria morphology in β-cells (Rieusset, 2018). It is possible that the increased Ca^2+^ concentration in the ER of TMEM24 KO cells causes a concomitant rise of Ca^2+^ in the mitochondria as these contact sites concentrate many of the key components of organellar Ca^2+^ homeostasis, including the MCU, the voltage-dependent anion channel (VDAC), SERCA, Presenilin-1 and IP3 receptors (Area-Gomez et al., 2009; Vecellio Reane et al., 2020). The MCU has low affinity for Ca^2+^ and is kept inactive under resting cytosolic Ca^2+^ concentrations (Marchi & Pinton, 2014). However, MCU is still important for maintaining basal energy production, likely via sensing Ca^2+^ microdomains formed by Ca^2+^ release from the ER at ER-mitochondria contacts (Rossi et al., 2019). Another intriguing possibility is that TMEM24 modulates the lipid composition of the mitochondria membranes, and that lack of this transport alters mitochondria function. TMEM24 has a strong preference for phosphatidylinositol, and the importance of this lipid for mitochondria function has been known since the 1960ies (Vignais et al., 1963). More recently, it has been shown that that the outer mitochondrial membrane is indeed rich in phosphatidylinositol (Zewe et al., 2020), and that phosphorylated derivatives of this lipid are required for mitochondrial function (Nagashima et al., 2020; Rosivatz & Woscholski, 2011). However, it remains to discover how phosphatidylinositol is delivered to the mitochondria. One possibility is that TMEM24 contributes to this transport and couples it to changes in Ca^2+^ concentration and energy demand. Interestingly, one study found that depletion or masking of PI(4,5)P_2_ on the mitochondria surface caused mitochondrial fragmentation, which could be prevented by PKC activation that would also trigger TMEM24 dissociation from the plasma membrane to enable interactions with the mitochondria (Rosivatz & Woscholski, 2011). A Ca^2+^-dependent feedback system to control mitochondria function may be particularly important in the β-cells, where mitochondrial metabolism is tightly coupled to Ca^2+^ influx in order to adjust insulin secretion and maintain blood glucose homeostasis.

## AUTHOR CONTRIBUTION

BX and SP performed all experiments and analyzed all data, except for the metabolic measurements presented in Fig. 4, which were done by CJ. BX and OI-H wrote the manuscript with input from SP, PG, CJ and PB.

## ACKNOWLEDGEMENT

We are grateful to Mrs. Antje Thonig for excellent help with molecular biology work and insulin secretion measurements and to Prof. Erik Gylfe and all members of the OI-H lab for input on the work. This study was funded by grants from The Novo-Nordisk Foundation, The Swedish Research Council, Åke Wibergs Stiftelse, Diabetesfonden and Exodiab (all to OI-H). P.G. is Research Director of the Fonds National de la Recherche Scientifique (Brussels).

## SUPPLEMENTARY FIGURES

**Supplementary Figure 1.**
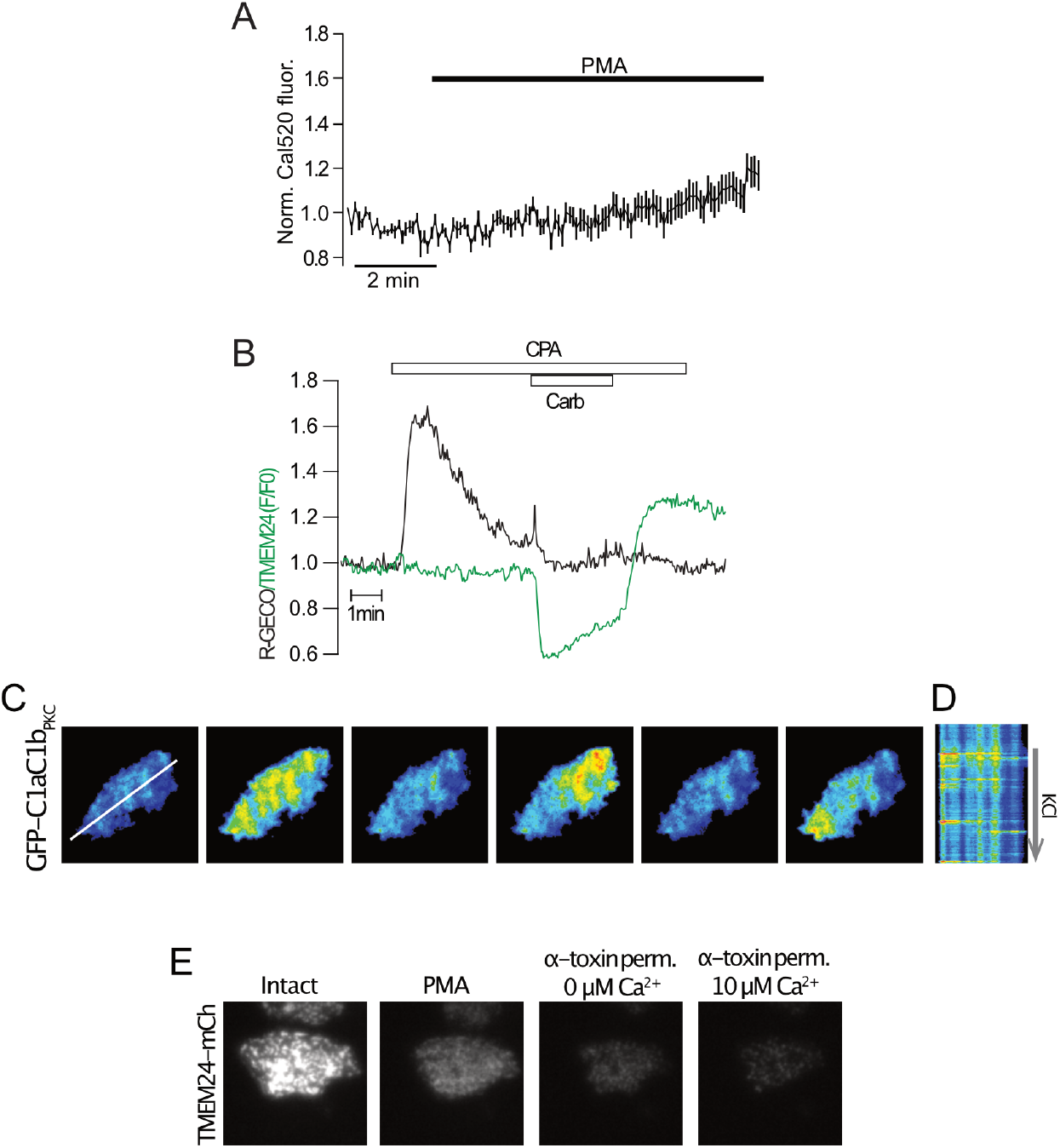
(A) Recordings of cytosolic Ca^2+^ in MIN6 cells loaded with the fluorescent Ca^2+^ indicator Cal520 following addition of 1 μM PMA. Data presented are means±SEM for 26 cells. (B) TIRF microscopy recording of R-GECO (black; Ca^2+^) and GFP-TMEM24 (green) fluorescence from a MIN6 cells exposed to 100 μM CPA and 10 μM carbachol. (C, D) TIRF microscopy images from a MIN6 cell expressing the DAG-sensor GFP-C1aC1bPKC during depolarization with 30 mM KCl. Notice the irregular formation of DAG at the plasma membrane, apparent as transient, localized fluorescence increase events in the kymograph in D. (E) TIRF microscopy images GFP-TMEM24 fluorescence from a MIN6 cell that is first exposed to 1 μM of the DAG analogue PMA, followed by permeabilization in an intracellular-like buffer and exposure to the indicated Ca^2+^ concentrations. Notice that most of the plasma-membrane-associated GFP-TMEM24 fluorescence is lost following addition of PMA.

## Notes

### Competing Interest Statement

The authors have declared no competing interest.

